# DYNAMIC CHANGES TO THE INTESTINAL ENVIRONMENT OCCUR THROUGHOUT RECOVERY FROM EXPERIMENTAL ISCHAEMIC STROKE

**DOI:** 10.1101/2025.04.25.650631

**Authors:** Rachel M. L. Martin, Isobel C. Mouat, Robert Whelan, Lizi M. Hegarty, Christopher J. Anderson, David H. Dockrell, Calum C. Bain, Gwo-Tzer Ho, Laura McCulloch

## Abstract

Stroke survivors experience a plethora of complications during recovery, including gastrointestinal symptoms. Intestinal dysfunction is reported to occur rapidly following stroke in both humans and animal models and alterations such as reduced barrier integrity, lymphocyte loss, and an altered microbiota have been suggested to contribute to poor neurological outcomes. Despite the persistence of gastrointestinal symptoms in many stroke survivors, how the intestinal environment changes over the course of stroke recovery remains poorly understood. Here, we use an experimental model of ischemic stroke to profile the gastrointestinal tract over a three month period of recovery.We have shown that experimental stroke leads to structural alterations to the colon, impaired transit times and an altered bacterial community composition. No impairments to barrier function were detected at any time point and transit times recover within 2 weeks post stroke. In contrast, structural and bacterial community alterations remain up to 3 months post stroke and are accompanied by abnormalities that develop only during chronic recovery, such as altered antibody coating of bacteria. These results suggest that the gastrointestinal system is dynamically altered over the course of experimental stroke recovery and that certain defects persist chronically after stroke.

## 1. INTRODUCTION

Stroke is a leading cause of death and disability with 143 million recorded prevalent cases of stroke worldwide in 2019 and an ever increasing prevalence (Feigin et al., 2021). Advances in hyperacute stroke care and the introduction of thrombectomy and thrombolysis for clot removal have improved survival rates. As a result, the number of stroke survivors is predicted to double by 2035 (King et al., 2020). Non-neurological complications remain an issue during recovery; infection and gastrointestinal dysfunction can impair neurological recovery and impact quality of life (Teh et al., 2018; Tuz et al., 2022). Gastrointestinal effects of stroke include dysphagia (Arnold et al., 2016), constipation (Li et al., 2017), and intestinal bleeding (Du et al., 2020; Fu, 2019). These complications often persist long-term and are linked to worsened clinical outcomes from stroke, including increased risk of recurrent stroke and mortality.

Intestinal dysfunction is well reported acutely after stroke, including gut barrier leakiness, reduced mucus production in animal models, and delayed transit time additionally in patients (Kumar et al., 2024; Li et al., 2017; Lim and Childs, 2013; Prame Kumar et al., 2023a; Stanley et al., 2016). The gut bacterial community, or microbiota, has been shown to be altered within 24 hours of stroke in both people and experimental animal models and dysbiosis has been independently associated with worsened functional and neurological outcomes (Benakis et al., 2020; Du et al., 2020; Fu, 2019; Rofes et al., 2018; Roth et al., 2020). Intestinal immune homeostasis is also compromised, (Conesa et al., 2023; Park et al., 2021) and loss of Peyer’s patch and mesenteric lymph node lymphocytes (Brea et al., 2021; Crapser et al., 2016; Schulte-Herbrüggen et al., 2009; Tuz et al., 2024) reported. Impairments to the mucus and epithelial cell barriers are thought to lead to translocation of bacteria, therein contributing to systemic inflammation (Houlden et al., 2016; Prame Kumar et al., 2023b). Additional factors that likely contribute to intestinal dysfunction during initial phases of recovery consist of reduced food consumption or nasogastric tube feeding, recumbency, and medications which alter gut transit or gastric pH, all of which can affect nutritional status and intestinal motility.

The established link between intestinal changes, particularly translocation of pathobionts, alongside the rapidly developing brain injury, has necessitated studies which have focused on hyperacute changes to the intestinal microenvironment. Intervention studies using antibiotic removal of the intestinal microflora or supplementation with short chain fatty acids after experimental stroke highlight the detrimental impact an altered bacterial community can have on the recovering brain (Benakis et al., 2020; Sadler et al., 2020). However, gastrointestinal issues often persist throughout recovery and are associated with risk of infection (Arnold et al., 2016), increased mortality (Fu, 2019; Rofes et al., 2018), and recurrent stroke (Du et al., 2020; Roth et al., 2020). Despite the chronicity of gastrointestinal complications, temporal characterisation of the functional, physiological, immunological and microbial changes to the intestine following stroke is currently lacking. Therefore, there is a limited understanding of the factors that may drive persistent intestinal complications after stroke.

In this study, we present foundational information into the impact of stroke on the intestine over the course of stroke recovery. Our findings reveal persistent macroscopic changes to colon structure up to three months post-stroke, despite a normalization of barrier function and transit time. Furthermore, we observe that the intestinal microbial community composition is altered during both acute and chronic recovery, with distinct shifts occurring in each phase, indicating that the post-stroke microbiota neither regains homeostasis nor is stable. Lastly, we find reduced antibody coating of bacteria, particularly during chronic stroke recovery, which appears to be independent of antibody availability and intestinal B cell numbers. We propose that prolonged and dynamic changes to intestinal homeostasis occur after stroke. Characterising the sequelae of events that occur in the intestine post-stroke provides the fundamental knowledge required for future experiments to determine mechanistic insight into the drivers of gastrointestinal dysfunction and the opportunity to uncover targeted interventions to improve intestinal health for survivors of stroke.

## 2. MATERIALS AND METHODS

### 2.1 Experimental stroke surgery

Experimental ischaemic stroke was achieved using the middle cerebral artery occlusion (MCAO) model. In brief, a 6-0 nylon monofilament with a 2-mm silicone coated tip (210 μm diameter; Doccol) was inserted into the external carotid artery and advanced through the internal carotid artery to occlude the middle cerebral artery. The filament was removed after 30 minutes to allow reperfusion, the neck wound was closed and animals recovered. Sham animals experienced the same surgical procedure, however, the filament was immediately retracted upon reaching the middle cerebral artery. Surgery was performed under isofluorane anaesthesia induced at 3.5% and maintained at 1.5% with O_2_ (200 ml/minutes) and N_2_O (400 ml/minutes). Core body temperature was maintained at 37 ± 0.5°C throughout the procedure with feedback controlled heating via a small animal physiological monitoring system (Harvard Aparatus) which also allowed monitoring of heart rate, respiration and blood oxygen saturation throughout the surgical procedure and animals were recovered in cages partially placed on a heated recovery mat for a minimum of 1 hour. Post-operative analgesia included application of lidocaine cream (LMX4 to the wound) and 0.1 mg/kg buprenorphine immediately after surgery and 24 hours post-surgery. Mice were recovered for five days, two weeks (12-15 days), or three months (61-95 days) following surgery. Post-stroke neurological deficits were measured over the course of recovery using the Clark focal neurological score (Clark et al., 1997).

### 2.2 Experimental design

All experiments used male C57BL/6 mice aged 8-12 weeks maintained under specific pathogen-free conditions and a standard 12-hour light/dark cycle with unlimited access to food and water. Mice were housed in individually ventilated cages of 4-6 animals per cage and were acclimatised for a minimum of one week before procedures. Experiments were carried out under a UK Home Office Project License in accordance with the @Animals (Scientific procedures) Act 1986 and Directive 2010/63/EU and were approved by the University of Edinburgh’s Animal Welfare and Ethics Review Board. The ARRIVE guidelines were consulted for experimental design, analysis, and reporting (Percie du Sert et al., 2020) and the IMPROVE guidelines were additionally used to maximise recovery after MCAO surgery to induce ischaemic stroke (Percie du Sert et al., 2017). Sample sizes were estimated based on previous data investigating immune changes during acute stroke recovery (McCulloch et al., 2017) using poweranalysis carried out in InVivoStat software (https://invivostat.co.uk/) to be sensitive to a 35% effect of experimental stroke at 80% statistical power at a 5% significance level. Animals were ear notched for identification and then randomized to experimental groups using Research Randomiser-an online random number generator (https://www.randomizer.org/). Animals were given an experimental identifier and scientists analysing experimental samples were blinded to experimental group. Data were unblinded upon completion of experiments. The average day two Clark focal neurological score was the same between timepoint groups. Sham and stroke mice were co-housed unless otherwise specified. Naïve controls were culled throughout experimental time course (5 days to 3 months) to provide a mixed age control group. Where appropriate, data from independent experiments were combined for analysis and are detailed in figure legends. Mice were euthanized by cardiac puncture while under isoflurane.

Post-hoc exclusion criteria were applied as follows: two naïve animals were excluded from analysis due to a cyst/tumor on abdomen or splenomegaly, and two stroke mice were excluded due to technical limitations of the procedure with failure to develop strokes, determined by a neurologic score <3 at day 1 of recovery and confirmed by histological analysis of brain. Exclusion of data from specific analyses are detailed in relevant methods sections, where appropriate.

### 2.3 Sample collections

Serially sampled faecal pellets were obtained with from the cage bottom while mice were individually caged, or colon contents were removed directly following cull and were snap frozen on dry ice and maintained at -80⁰C until analysis. Within an experiment, faecal pellets were collected within the same three-hour period of the day, the timing of which was consistent between collection days. Upon cull, the length of the colon was measured. Colon samples taken for histology were flushed with phosphate-buffered saline (PBS), cut longitudinally, rolled into Swiss roll shape, frozen on dry ice, and stored at -80°C. Colon samples for flow cytometry were cut open longitudinally, washed in PBS, cut into approximately one centimeter pieces, and placed into PBS on ice. Four to eight Peyer’s patches were removed from each animal and placed into PBS on ice.

### 2.4 16S rRNA sequencing

To examine the composition of the bacterial microbiota, 16S rRNA sequencing was performed. DNA was extracted from 40-200 mg stool by the NU-OMICS DNA Sequencing facility (Northumbria, UK). Bacterial DNA was isolated with the QIAGEN PowerLyzer PowerSoil kit (Qiagen) following manufacturer’s instructions. Library preparation was carried out by NU-OMICS (Northumbria University) based on the Schloss wet-lab MiSeq SOP (Kozich et al., 2013). Briefly, PCR was performed using the 1x KAPA2G Robust HotStart ReadyMix, 0.5 µM each primer and 1 µl of template DNA with the following cycling conditions: 95⁰C 2 minutes, 30 cycles 95⁰C 20 seconds, 55⁰C 15 seconds, 72⁰C 5 minutes with a final extension 72⁰C 10 minutes. A negative and positive (Zymobiomics Microbial Mock community DNA standard) control sample were included in each 96 well plate and carried through to sequencing. PCR products were quantified with the Quant-iT™ PicoGreen™ dsDNA Assay (Invitrogen) and each sample was normalised to 10nM and then pooled within each 96-well plate. Each pool was quantified using fragment size determined by BioAnalyzer (Agilent Technologies) and concentration by Qubit (Invitrogen) and pools were combined in equimolar amounts to create a single library. The library was then denatured using 0.2N NaOH for five minutes and diluted to a final concentration of 4.5 pM, supplemented with 20% PhiX and loaded onto a MiSeq V2 500 cycle cartridge.

### 2.5 16S rRNA bioinformatics analysis

DADA2 was implemented using the Nextflow nf-core ampliseq pipeline (version 2.5) for the 16S rDNA bioinformatic analysis (Straub et al., 2020). Paired-end sequences were filtered and trimmed with -- trunclenf 220 and --trunclenf 170 and default parameters. Taxonomy was assigned using the Silva 138.1 reference database (Quast et al., 2012). The phyloseq package (version 1.38.0) was used to generate a phyloseq object from the DADA2 output for downstream analysis (McMurdie and Holmes, 2013). The amplicon sequence variant (ASV) abundance table was normalised using the total sum scaling (TSS) method to relative abundances, and low abundant ASVs were filtered out (<0.01%). Alpha diversity analyses were generated using phyloseq (version 1.38.0) and the Shannon diversity index was used. Analyses was performed on both raw unfiltered and rarefied ASV-level. Beta diversity analyses was performed with Vegan (10.32614/CRAN.package.vegan), using Bray-Curtis dissimilarity matrices from TSS normalised and arcsine square root transformed data.

### 2.6 Histological analysis

Colon Swiss rolls were embedded in optimal cutting temperature compound (Tissue-Tek) and 6 μm sections were cut on an HM525 NX Cyrostat (Thermo Fisher Scientific) at -20⁰C. Sections were stored at -20°C until further analysis. Sections of fresh frozen colons were hematoxylin and eosin (H&E) stained. Briefly, sections were defrosted then stained with haematoxylin for five minutes, washed in distilled water, immersed in Scott’s tap water (CellPath) for 30 seconds followed by a 30 second wash in distilled water. Sections were then stained with eosin for three minutes then washed with distilled water. Sections were dehydrated by soaking for two minutes in 70% > 80% > 95% > 100% ethanol, fixed for five minutes in xylene, then coverslips were applied using DPX mountant (Merck). To visualise the mucus barrier, cross sections of colon, with faecal content intact, were dissected into Carnoy’s fixative for 3 hours before removal into 70% ethanol, processed and embedded in parrafin blocks. Sections of 6 μm were cut and deparrafinised by soaking for ten minuntes in xylene, followed by rehydration in decreasing concentrations of ethanol 100%>95%>80%>70% for five minutes each. Sections were stained in hematoxylin for five minutes, washed in distilled water, stained for 30 minutes in Alcian Blue (pH 3.0), washed in distilled water and then stained with eosin for 3 minutes. Sections were then dehydrated and coverslipped as above.

### 2.7 Immunofluorescence

Fresh frozen 6 μm colon sections were fixed in ice-cold acetone for ten minutes, washed in 0.05% bovine serum albumin in PBS (PBS-BSA) and blocked using normal goat serum (Jackson ImmunoResearch). Primary antibodies were incubated for one hour at room temperature or overnight at 4 °C. Tight junctions (ZO-1), epithelial cells (EpCAM), and proliferating cells (Ki-67) detected using the antibodies described in **Table 1**. Mucin producing cells were detected using the lectin Ulex Europaeus Agglutinin I (UEA I) directly conjugated to rhodamine (1:200; Vector Laboratories; RL-1062-2). Following addition of primary antibody, sections were washed in PBS-BSA and where relevant, species specific secondary antibody, or streptavidin, conjugated to Alexa Fluor (AF) dyes, decribed in **Table 2**, were added to sections and incubated for one hour. Sections were washed again and DAPI was added (1:5000; Thermo Fisher Scientific) for ten minutes. Sections were given a final wash in PBS-BSA and mounted in Dako Fkuorescence Mounting Medium (Agilent, S3023). Control sections were incubated only with secondary antibody and/or species specific normal serum.

**Table 1.**
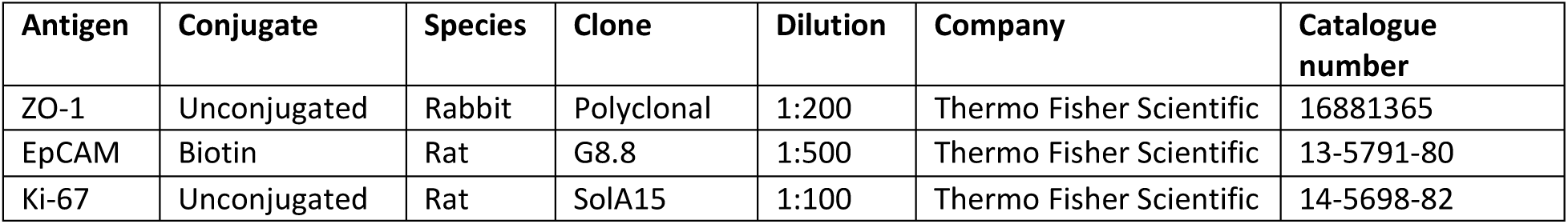
Primary antibodies for immunostaining.

**Table 2.** Secondary detection for immunostaining.

| Secondary detection | Conjugate | Species | Dilution | Company | Catalogue number |
| --- | --- | --- | --- | --- | --- |
| Anti-rabbit IgG | Alexa Fluor 647 | Goat | 1:500 | Thermo Fisher Scientific | A21245 |
| Anti-rat IgG | Alexa Fluor 555 | Goat | 1:200 | Abcam | ab150166 |
| Streptavidin | Alexa Fluor 488 | N/A | 1:500 | Thermo Fisher Scientific | S32354 |
| Streptavidin | Alexa Fluor 647 | N/A | 1:500 | Thermo Fisher Scientific | S32357 |

Apoptotic TUNEL^+^ cells were measured using the ApopTag Fluorescein Direct In Situ Apoptosis Detection Kit (Merck, s7110) to detect DNA fragmentation according to the manufacturers protocol. Sections were fixed in 1% paraformaldehyde in PBS, washed in PBS and post-fixed in ice-cold ethanol: acetic acid (2:1). DNase treated positive controls provided in the kit were used to confirm staining. Sections were incubated in reaction mix containing DIG-labelled dUTP and terminal deoxynucleotidyl transferase for 60 minutes. After washing, fluorescein-conjugated anti-DIG was applied to sample, which was then incubated in the dark for 30 minutes. Following TUNEL staining, immunolabelling for proliferating (Ki-67) and epithelial (EpCAM) cells was performed as described above. Sections were then washed and mounted in Dako fluorescent mounting media (Agilent, S3023).

### 2.8 Image analysis

Images of colon Swiss rolls were taken on Axio Scan.Z1 (Zeiss). Whole tissue was scanned at 20x magnification. A single five day sham animal was excluded from all image analysis due to poor quality of tissue. Files were exported as .czi for use in QuPath software (version 0.4.3).

Crypt depth was measured using the measure tool and drawing a line from the base to the tip of the crypt. The average crypt depth was recorded by measuring ten intact crypts across the entirety of the colon section per animal. Muscularis thickness was quantified in QuPath by drawing a line across the muscularis at 12-18 locations along the length of the colon. Nuclear density was assessed by generating five regions of interest measuring 200 x 200 μm which were placed randomly along sections of the crypt or muscularis. The number of nuclei in each region of interest was determined using the cell detection tool within QuPath.

Ki-67, UEA-1 and ZO-1 analyses were carried out on regions of interest where the colon structure was clearly identifiable, any folded or damaged tissue was not included in analysis. To quantify Ki-67^+^ proliferating cells, UEA-1^+^ mucin-producing cells and ZO-1^+^ tight junctions the epithelial and lamina propria layer of tissue was selected using the brush tool in Qupath. Positive cells were detected and counted using the positive cell detection tool in QuPath and normalised to the area analysed to provide UAE-1^+^ and ZO-1^+^ cells per mm^2^ for each animal.

### 2.9 Faecal supernatant and protein assays

Faecal pellets were suspended in PBS at a concentration of 100 mg faeces per 1 ml of PBS. Following 20 minutes of incubation on ice, samples were homogenized 2 x 30 seconds at 6500 rpm (Precelleys homogenizer) and centrifuged at 8,000 x g for 10 minutes to pellet the bacteria. Supernatant was recovered and the bacterial pellet and supernatant were each snap frozen and stored at -80°C until analysis. Total protein in faecal supernatants was measured using the Pierce BCA Protein Assay Kit (ThermoFisher Scientific, A55865). Lipopolysaccharides (LPS) in plasma was measured with the Pierce Chromogenic Endotoxin Quant Kit (ThermoFisher Scientific, A39553) from samples diluted 1:75 and run according to manufacturer’s instructions. Calprotectin was measured in faecal supernatant diluted 1:2 with the Mouse Calprotectin ELISA Kit (Abcam, ab263885). To measure the abundance of various cytokines and antibodies in faecal supernatant, respectively, the LEGENDplex™ MU Th Cytokine Panel (Biolegend, 741044) and LEGENDplex™ Mouse Immunoglobulin Isotyping Panel (Biolegend, 740493) kits were used and analysed on an Attune NxT Flow Cytometer (Thermo Fisher Scientific). All procedures were carried out in accordance with manufacturer’s instructions.

### 2.10 Gut transit time

Transit time experiments were based on previously published methods (Schonkeren et al., 2023). As recommended, experiments were carried out at night under red light during the animals’ awake cycle. Mice were individually housed, fasted for 1-2 hours, and orally gavaged with 200 μl FITC-Dextran 70 (Sigma, 46945) prepared at 50 mg/ml in PBS. Upon gavage, access to a pre-weighed amount of food (normal chow) was returned. Cages were examined every three minutes using UV torches and the time of the first fluorescent pellet was recorded. The amount of food eaten (weight of initial minus remaining food) and faecal pellets produced was measured over the course of the experiment. Food weight data was not collected at first experimental collection and therefore data on food eaten per hour is missing for some animals (n=2 sham; n= 3 stroke) at the five day timepoint.

### 2.11 Isolation of intestinal leukocytes

To isolate leukocytes from the colon, tissue was cut into 1 cm pieces, washed three times with 10 ml ice cold FACS buffer (PBS with 2% FCS and 1mM EDTA (VWR, R013)), and intra-epithelial leukocytes were isolated by incubating at 37°C for 30 minutes in prewarmed stripping buffer (RPMI-1640 with 5% FCS, 1mM DTT (Thermo Fisher Scientific, R0861), and 1 mM EDTA) while shaking at 200 rpm. Intra-epithelial leukocytes were then placed through a 100 μm sieve, washed with 10 ml ice cold FACS buffer, and placed on ice until staining. To isolate lamina propria leukocytes, the remaining tissue was washed with 10 ml ice cold FACS buffer and incubated at 37°C for 45 minutes in prewarmed digestion buffer (RPMI-1640 with 1 mg/ml Collagenase D (Merck, 11088858001), and 20 μg/ml DNase1 (Roche, 101104159001)) while shaking at 200 rpm; the tissue and supernatant was then passed through a 70 μm sieve and washed with 10 ml ice cold FACS buffer. Peyer’s patches were passed through a 70 μm sieve to generate single cell suspension and washed with 10 ml ice cold FACS buffer.

### 2.10 Flow cytometry

Proportions of Peyer’s patch, intraepithelial, and lamina propria lymphocytes were quantified by flow cytometry. Cells were incubated with FC block (purified anti-mouse CD16/32, Biolegend, 101301), stained at 4°C in the dark with the antibodies described in Table 3, stained with 0.1 μg/ml DAPI (4′,6-diamidino-2-phenylindole, Thermo Fisher Scientific, D3571), and analysed with an LSRFortessa II Cytometer (BD Biosciences). Single-stain controls were prepared using UltraComp eBeads (Thermo Fisher Scientific) and counting beads were 123count eBeads™ Counting Beads (Thermo Fisher Scientific). Data were analyzed using FlowJo (V10).

**Table 3.**
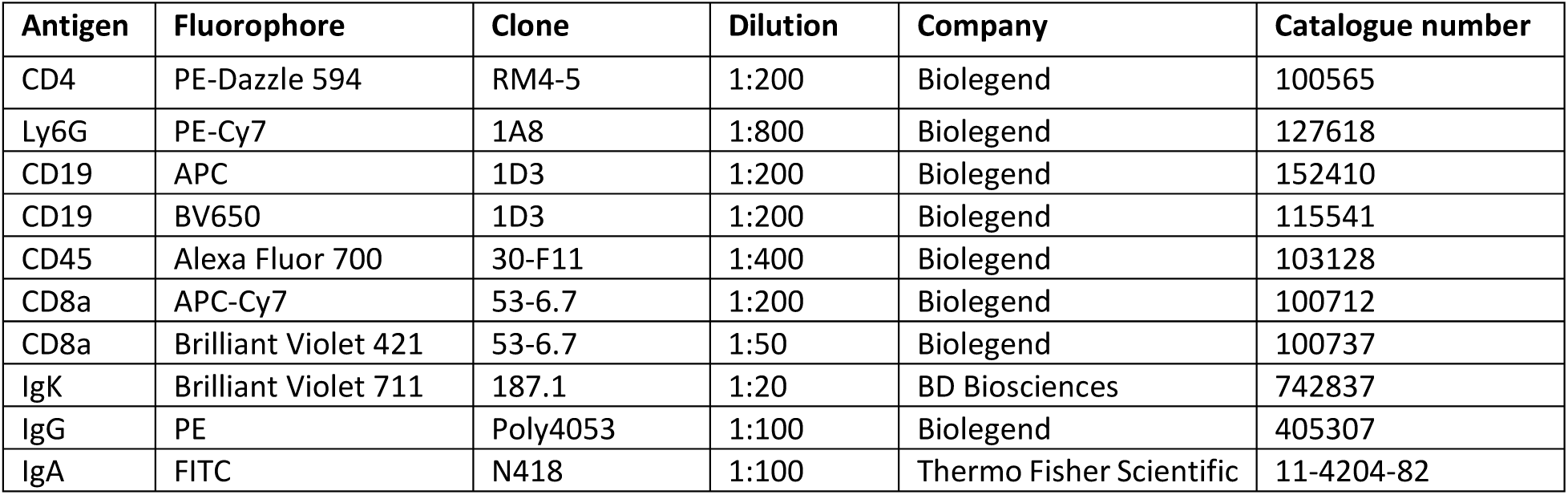
Flow cytometry antibodies.

### 2.11 Antibody coating of faecal bacteria

Bacterial pellets collected during preparation of faecal supernatant were resuspended in 1 ml FACS buffer and filtered through 70 μm sieve. Samples were stained with 17 μM SYTO-60 (Thermo Fisher Scientific, S11342) for 30 minutes at 4°C. Next, samples were stained with antibodies against IgK, IgG and IgA (**Table 3**) for 30 minutes at 4°C and subsequently with AF488-streptavidin at 1:500 (Thermo Fisher Scientific, S32354) for 60 minutes at 4°C. Data was aquired on a NovoCyte cytometer (Agilent) and data were analyzed using FlowJo (V10).

### 2.12 Statistical analyses

With the exception of 16s rRNA bioinformatic analysis, data presentation and statistical analyses were performed using GraphPad Prism software (version 9.5.1, GraphPad Software Inc.). For normally distributed data, differences were tested using one -way or analysis of variance (ANOVA) with Dunnett’s multiple comparisons test, comparison to naïve or stroke. Data that was not normally distributed according to Shapiro-Wilk was either analysed by Kruskal-Wallis or was log transformed and analysed by one-way ANOVA, as indicated in figure legend. Data with unequal standard deviations, determined by the Brown-Forsythe test, were analysed by Brown-Forsythe and Welch ANOVA tests. Ordinal data were analysed by Kruskal-Wallis. For data with two independent variables and with different numbers of observations in each group were analysed by fitting a mixed-effects model (REML) with the Geisser-Greenhouse correction. Multivariate analysis was performed by permutational multivariate ANOVA (PERMANOVA). Results are presented as mean ± SD unless stated otherwise. P-values indicated by asterisks as follows: ****p < 0.0001, ***p < 0.001, **p < 0.01, *p < 0.05.

## 3. RESULTS

### 3.1 Changes to colon morphology persist across stroke recovery

To investigate the intestinal microenvironment over the course of stroke recovery, animals underwent experimental stroke or sham surgery and were recovered to five days, two weeks or three months. Substantial neurological deficits were observed acutely after stroke with gradual recovery thereafter (**Fig. 1A**). By three months, animals had nearly recovered to baseline, though mild neurological symptoms persisted, including one-sided torso flexion and assymetrical whisker reflexes. Stroke animals experienced up to 20% loss in body weight acutely and never fully regained weight to the levels seen in sham controls (**Fig. 1B**). Unexpectedly, during the mid-recovery phase (5d-2w), average weight gain was similar between sham and stroke animals. However, during the chronic phase of recovery (2w-3m), stroke animals exhibited reduced weight gain compared to sham controls (**Fig. 1C**).

**Figure 1:**
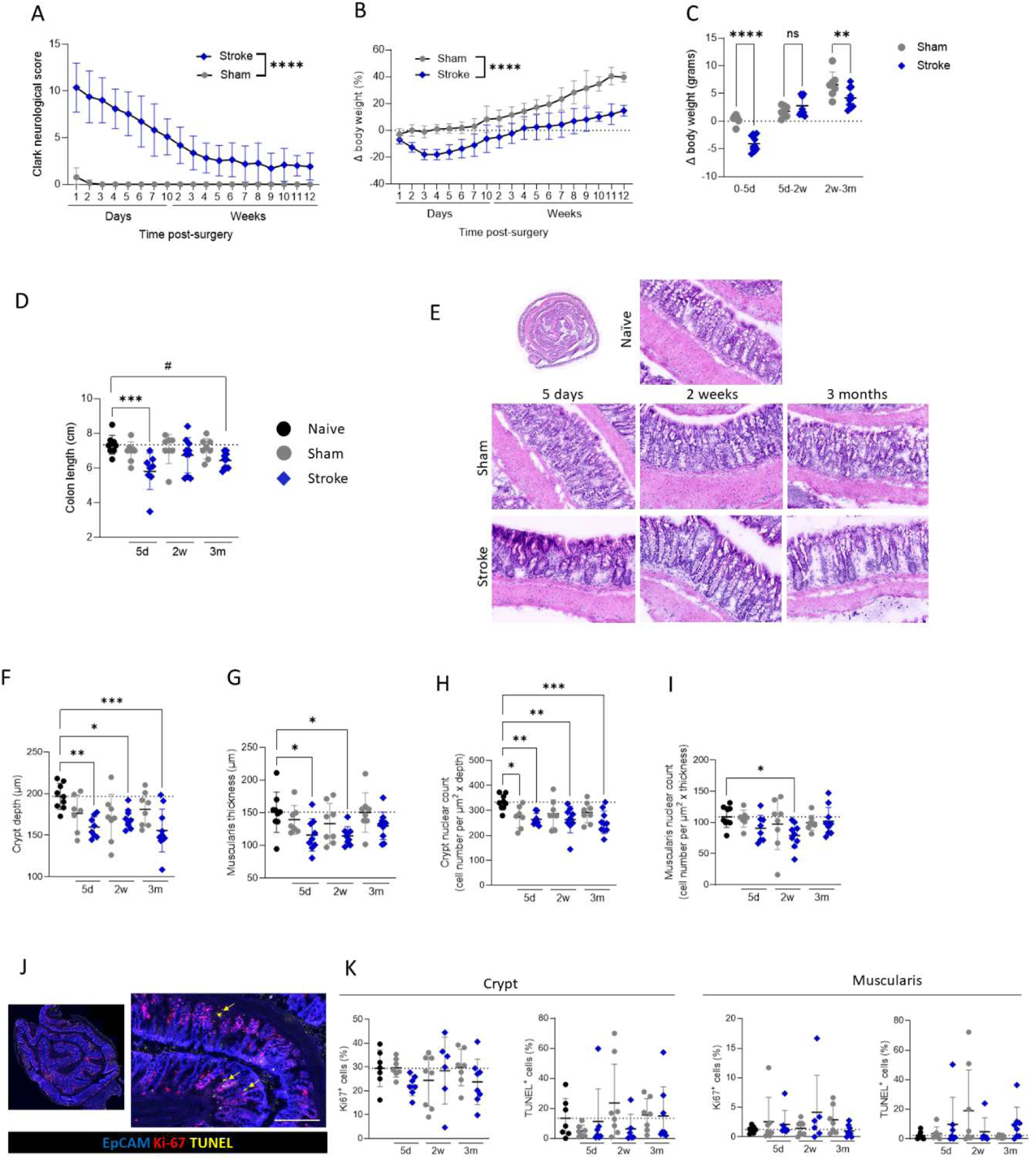
Changes to colon morphology during acute and chronic stroke recovery. Sham control (grey circles) and MCAO stroke (blue diamonds) animals were recovered for up to 3 months following surgery. **(A)** Clark focal neurological score and **(B)** body weight change measured over recovery. **(C)** Body weight change (in grams) during phases of recovery: 0-5 days, 5 days – 2 weeks, 2 weeks – 3 months. **(D)** Colon length measured upon cull (#, p=0.052). **(E)** Representative images of Haematoxylin and Eosin (H&E) stained colons. **(F)** Measurement of crypt depth and **(G)** muscularis thickness. **(H)** Crypt nuclear count normalised to crypt depth, and **(I)** muscularis nuclear count normalized to thickness in H&E stained colons. **(J)** Representative images of immunostaining for epithelial cells (EpCAM; blue), proliferation (KI-67; red) and cell death (TUNEL; yellow; arrows) scale bar 200 μm. **(K)** Measurement of the percentage of KI-67+ or TUNEL positive cells in the crypt and the muscularis. Analysed by mixed-effects model with the Geisser-Greenhouse correction **(A, B)**, multiple unpaired t-tests **(C, K)**, or one-way ANOVA with Dunnett’s multiple comparisons test with comparison to naïve **(D, F-I)**. N=7-10 per group. Data shown as mean ± S.D. Dotted line demarks the mean of the naïve group.

In certain murine models of intestinal inflammation, colon length is used often used as a crude but reliable proxy measure of colonic inflammation (Liang et al., 2024; Seo et al., 2024). Of note, stroke led to a significant reduction in colon length (approximately 25% shorter than those of controls). Surprisingly, this colon shortening was also evident at three months post-stroke (**Fig. 1D**), indicative of persistent changes to the colonic environment/structure. To assess macroscopic structure, H&E staining was performed on sections from colon Swiss rolls (**Fig. 1E**). Crypt blunting, indicated by a reduced crypt depth, was observed at all timepoints post-stroke (**Fig. 1F**). Additionally, thinning of the muscularis in the colon was noted at five days and two weeks post-stroke, although muscularis thickness had returned to baseline levels by three months post stroke (**Fig. 1G**). Cell numbers were reduced in the crypt at all timepoints (**Fig. 1H**), however a reduced cellularity was also observed at 5 days post sham surgery suggesting at this time point, the impact of recovering from surgery may play a role. Reduced cellularity was observed in the muscularis layer only at two weeks post-stroke (**Fig. 1I**). Despite structural changes to the colon, no clinical or histological signs of colitis were detected at any time point.

To determine if the macroscopic changes were associated with altered cell turnover, we performed immunofluorescent staining of apoptosis (TUNEL) and proliferation (marked by Ki-67 expression) within the colonic crypts and muscularis (**Fig. 1J**). There was a trend of reduced Ki67^+^ cells within the crypt at five days post-stroke compared with naïve and sham groups, but otherwise there was no evidence of altered cell proliferation or apoptosis at any timepoint measured (Fig. 1K). As cell density within these structures is unchanged (**Fig. S1**), atrophy of the crypt and muscularis may be driving the observed reduction in total cell number.

Together these findings suggest persistent alterations to intestinal morphology during chronic stroke recovery, warranting further investigation into the intestinal environment and functional capacities throughout stroke recovery.

### 3.2 Stroke-induced functional impairments in the gut recover at chronic timepoints

Next, selected intestinal functions were assessed throughout stroke recovery, including barrier integrity, mucus production, and transit time. First, to investigate gut barrier integrity we examined expression of the tight junction protein ZO-1 by colonic epithelial cells, which has been reported to display reduced expression in the colon in the first 24 hours after experimental stroke (Wen et al., 2019). In agreement with studies showing resolution of barrier integrity 24 hours post experimental stroke (Prame Kumar et al., 2023a), no changes in ZO-1 expression were detected in the colonic epithelium at any of the recovery time points assessed (Fig. 2A, B). Additionally, total protein content in faecal supernatant was unchanged (Fig. 2C), suggesting the outward intestinal barrier is intact at these recovery time points. Inward barrier integrity was examined by measuring bacterial LPS in plasma; only two animals at five days after stroke had detectable levels of circulating LPS, suggesting that the inward gut barrier is also largely intact at the time points assessed (Fig. 2D). Next, to examine the presence of goblet cells, we assessed staining with UAE-1, a lectin that binds to fucosyl glycoconjugates on goblet cells. This showed that UAE-1 staining was reduced in both sham- and stroke-operated animals at the five day recovery time point. UAE-1 levels returned to baseline by two weeks and three months post-stroke (Fig. 2E, F). That these effects were observed in both sham-operated and stroke groups suggests that the number of goblet cells and/or mucin production is likely an effect of surgery. The mucus barrier was further assessed in sections of transverse colon, with intact faecal material, stained with alcian blue from animals 5 days (Fig. 2G). In support of UAE-1 quantification, mucus barrier thickness was reduced in animals 5 days post-stroke in comparison to naïve controls, with sham animals demonstrating an intermediate phenotype (Fig. 2H).

**Figure 2:**
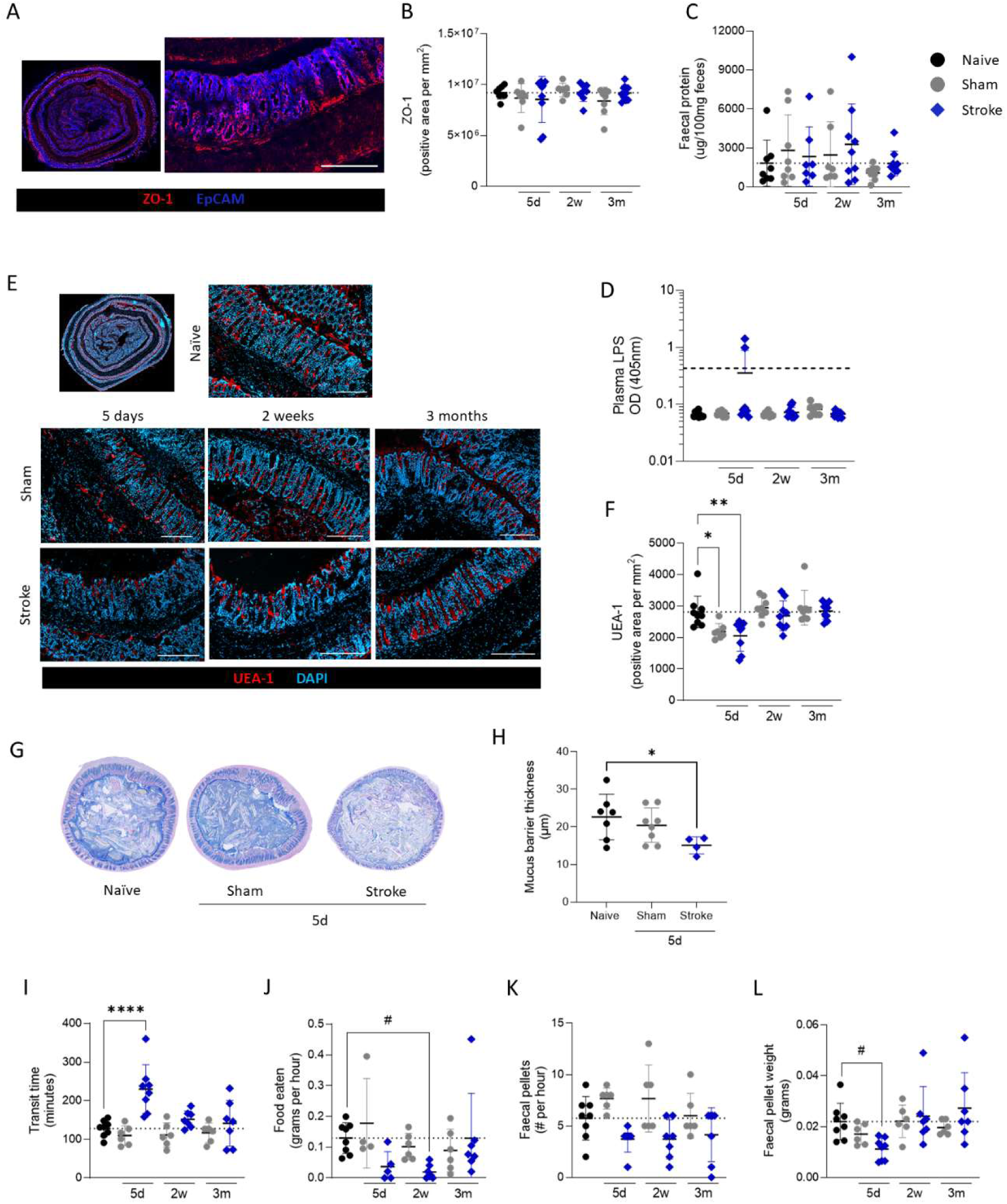
Selected intestinal functional impairments in acute stroke recovery at chronic timepoints. Colons and plasma were collected from MCAO stroke (blue diamonds) and sham control (grey circles) mice at 5 days, 2 weeks, and 3 months of recovery. **(A)** Colons were immunostained for tight junctions (ZO-1; red) and epithelial cells (EpCAM; blue); scalebar 200 μm **(B)** Quantification of ZO-1+ tight junctions in colonic crypts. **(C)** Faecal protein concentration was measured in faecal supernatant. **(D)** LPS quantified in plasma. **(E)** Representative images of colons immunostained for UEA-1 (red) and DAPI (blue) and associated **(F)** quantification of UEA-1. **(G)** Alcian blue and haematoxylin staining of colon cross-section with faecal material intact and associated (**H**) quantification of mucus barrier thickness. **(I)** Time (in minutes) reported for orally gavaged FITC-Dextran to appear in faecal pellet. **(J)** Amount of food (grams per hour) eaten (#, p=0.076) and **(K)** faecal pellets produced over the course of an hour. Random exlcusion of a subset of animals due to technical contstraints described in methods. **(L)** Weight of individual faecal pellet (#, p=0.060). Scalebar 200 μm **(A, E)** Data presented as mean ± SD. Dotted line is the mean of the naïve group **(B, C, F, I-L)** or the limit of detection **(D)**. N=7-9 per group **(B-D, F)**, n=4-8 **(H)**, n=6-8 **(I-L)**. Analysed by one-way ANOVA with comparison to naïve **(B-D, F, H, J-L)** or Kruskal-Wallis test **(I)**.

Gut transit time was assessed by measuring the time taken for gavaged FITC-Dextran to appear in faeces, and was significantly delayed at five days post-stroke which normalised at later recovery time points (Fig. 2I). Food intake was measured throughout the experiment and stroke animals showed a tendancy to consume less food at five days of recovery, which may contribute to the acute delayed transit time and reduced weight of stroked animals (Fig. 2J). However at 2 weeks post-stroke, food intake remains reduced but transit time and loss of body weight are similar to controls, suggesting additional factors may contribute to these phenotypes (Fig. 1C). In agreement with data on food intake, the number of faecal pellets produced per hour (Fig. 2K) and the faecal weight of pellets produced (Fig. 2L) were also reduced early after stroke.

Together these data demonstrate that certain changes to gut functions occur acutely after stroke and do not persist into chronic recovery. Furthermore effects on goblet cell number and mucus barrier may in part be driven by surgery. Additionally, changes to mucus barrier functions, food intake and gut transit time may contribute to intestinal dysfunction early after stroke but are unlikely to contribute to any persistant alterations to intestinal homeostasis.

### 3.3 Stroke induces dynamic changes to the microbiome which persist into chronic recovery

Shifts in the composition of the intestinal microbiota is well reported during acute stroke recovery in both animal models and human studies (Singh et al., 2016; Xu et al., 2021). To profile the community composition induced by our model of stroke over the course of recovery, we performed 16S rRNA sequencing of faecal material from naïve and stroke animals. In these experiments, stroked animals were co-housed with sham-operated controls to aid the welfare and recovery of their stroked cagemates (Percie du Sert et al., 2017). Co-housing resulted in an overlapping community composition between treatment groups (**Fig. S2A**). However, when animals were separately housed, the microbiome of naïve and shams were distinct from that of stroke animals (**Fig. S2B**). These results suggest that the microbiota of stroke animals can seed their sham operated cagemates when they are co-housed for welfare reasons. For this reason, microbiota abundances from pre-stroke and separately housed animals were used as a baseline comparison for stroke-induced changes to bacterial community composition in the above analyses.

The microbial community composition was significantly altered between naïve and post-stroke sample groups at all timepoints; interestingly, community composition was different between early (five day and two week) and late (three month) recovery timepoints (Fig. 3A). These results suggest that stroke affects the intestinal microbiota throughout recovery in a dynamic manner, with distinct compositions observed during acute versus chronic recovery.

**Figure 3:**
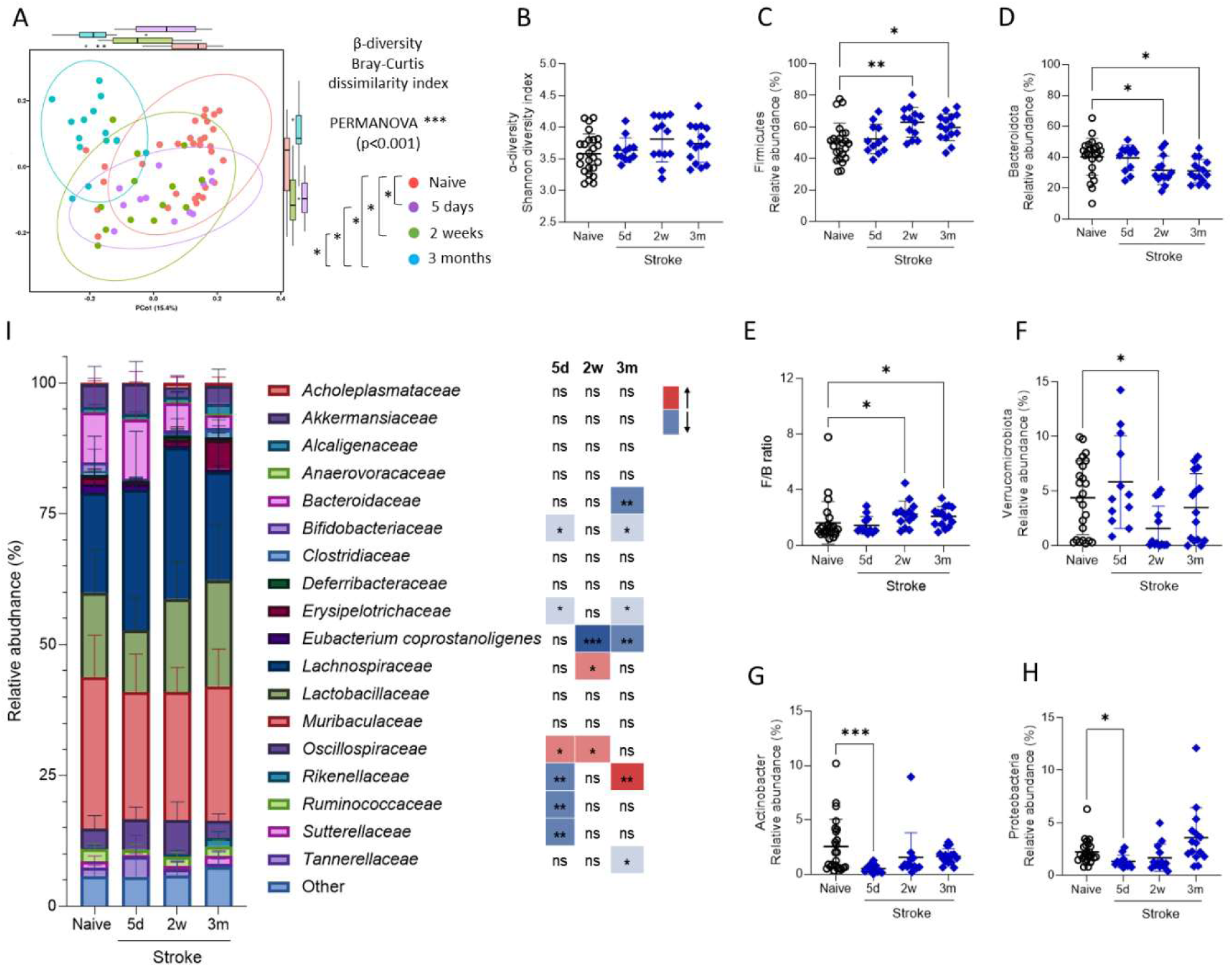
Stroke induces dynamic changes to the microbiome which persist into chronic recovery. Sham control (grey circles) and MCAO stroke (blue diamonds) animals were recovered for up to 3 months following surgery and 16S rRNA sequencing was performed on faecal material. **(A)** Bray-Curtis dissimilarity index, analysis during stroke recovery and **(B)** Shannon diversity index. **(C-H)** Relative abundance of various phyla, including Firmicutes, Bacteroidota, Verrucomicrobiota, Actinobacter, and Proteobacteria and the ratio of Firmicutes to Bacteroidota. **(I)** Relative abundance of bacterial families with an abundance of >1% of total population. Bacterial families comprising < 1% were included in the group defined as other. Data presented as mean ± SD **(A, C-I)** or 95% confidence internal **(B)**. Data are pooled from two independent experiments, n=12-25. Naïve (open circles) are samples from mice without sham or stroke surgery and housed separately from post-surgery mice. Data shown as mean ± SD. Data analysed by one-way ANOVA, or Kruskal-Wallis test if SD’s significantly different by the Brown-Forsythe test, with comparison to naïve **(B-H)**, PERMANOVA, with and without multiple comparisons test **(A)**, or mixed-effects analysis with Dunnett’s multiple comparisons test with comparison to baseline **(I)**.

The diversity within the microbial community of individual samples was measured with the Shannon diversity index and shown to be unchanged throughout stroke recovery (Fig. 3B), demonstrating that the observed dissimilarity between sample groups is not due to altered total species evenness and richness. Therefore, we began investigating the relative abundance of various bacterial species.

Firmicutes and Bacteroidetes are the two most dominant phyla of the mouse gut microbiota (Beresford-Jones et al., 2022) and the ratio of these has been linked to various conditions in humans including obesity (Ley et al., 2006), cardiovascular disease (Huang et al., 2024), and sepsis (Sankar et al., 2024). In this model of ischaemic stroke, significant increases in the abundance of Firmicutes (Fig. 3C) and reductions in Bacteroidota (Fig. 3D) were detected at two weeks and three months after stroke resulting in altered Firmicutes/Bacteroidota ratios at these time points in comparison to naïve controls (Fig. 3E). Further changes detected at the phylum level included reduced abundance of Verrucomicrobiota at two weeks (Fig. 3F), and reduced Actinobacter (Fig. 3G) and Proteobacteria (Fig. 3H) at five days post-stroke, in comparison to naive controls.

Investigation of gut microbiome at the family level further demonstrated different temporal trajectories. The abundance of some families showed changes only at five days (*Ruminococcacea, Sutterellaceae)* or three months (*Bacteroidaceae, Tanerellaceae*) post-stroke. Other families showed opposing changes in abundance at five day and three month recovery timepoints (*Rikenellaceae*) and some showed changes at five day time points which returned to baseline at two weeks and reoccurred at three months (*Bifidobacteriaceae, Erysipelotrichaseae*; Fig. 3I).

In summary, the intestinal microbiome displayed dynamic changes throughout stroke recovery. Distinct microbial alterations during acute and chronic timepoints suggest that different factors may be responsible for driving microbiota changes across stages of recovery. Reduced eating (Fig. 2J), impaired transit (Fig. 2I) and reduced mucus barrier (Fig. 2H) may contribute to population changes measured during acute stroke recovery but again these are unlikely to be associated with chronic microbial alterations.

### 3.4 Disrupted intestinal immune homeostasis occurs throughout stroke recovery

Intestinal homeostasis is crucial for normal physiological and barrier functioning and is tightly regulated by a number of host response mechanisms. Shifts towards either inflammation or immunosuppression can disrupt intestinal morphology, functionality, and bacterial community composition. Following stroke, immune homeostasis is disrupted in various peripheral tissues, with a combination of hyporesponsiveness and inflammation observed (Endres et al., 2022). Given this, we next measured soluble immune factors in the intestinal lumen over recovery.

Reductions in colonic luminal concentrations of cytokines IL-4, IL-9, and IL-22 were detected five days after stroke and a similar trend was observed for IL-13 (Fig. 4A**-D**). In contrast, concentration of IFNγ, TNF, IL-2, IL-17F, IL-5, IL-6, IL-17A, and IL-10 (**Fig. S3A**) remained relatively unchanged throughout stroke recovery. No increases in luminal cytokine concentrations were detected in the faecal supernatant at any time point.

**Figure 4:**
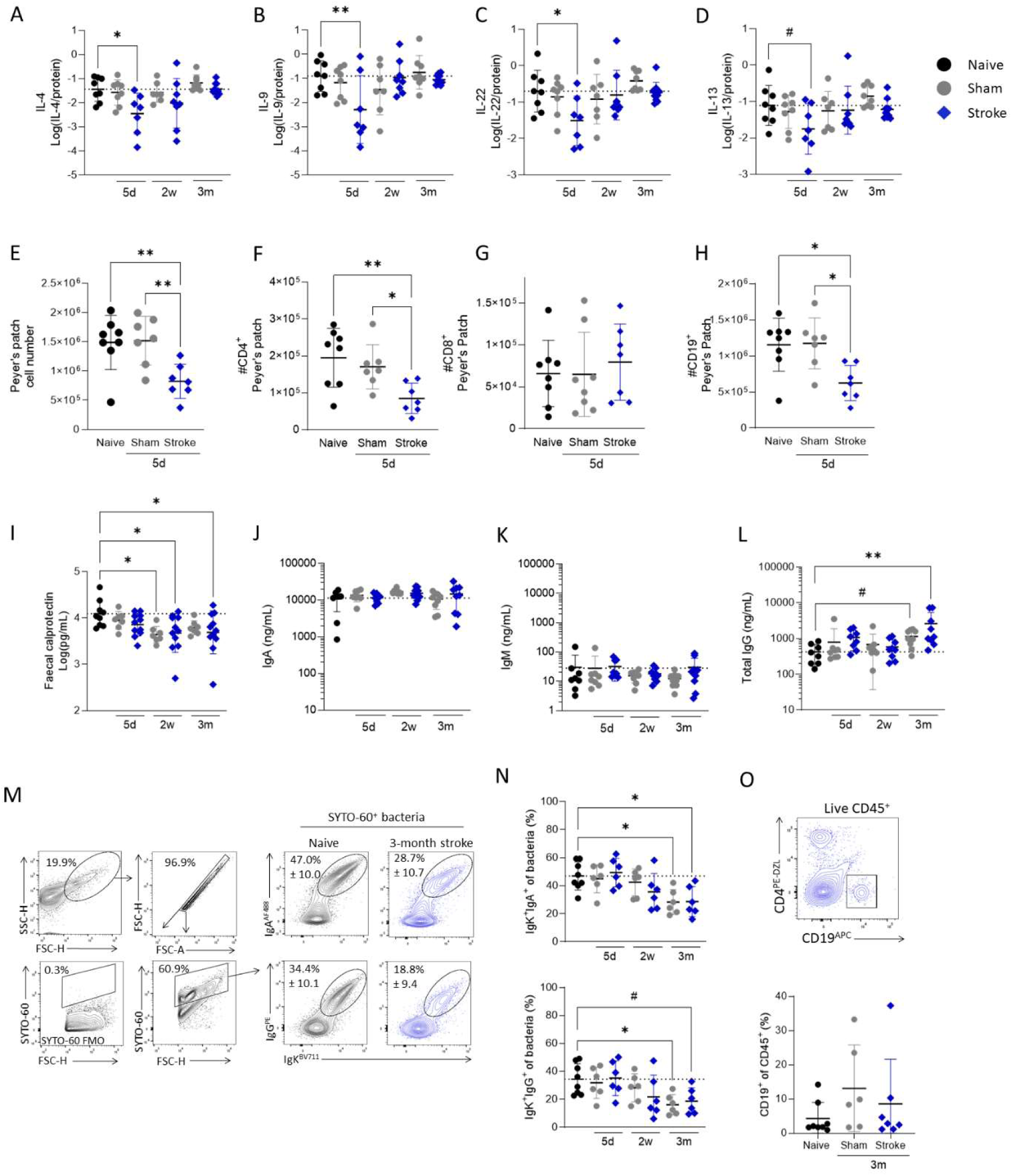
Disrupted intestinal immune homeostasis throughout stroke recovery. Faecal samples were processed for faecal supernatant and a bacterial fraction from sham control (grey circles) and MCAO stroke (blue diamonds) animals. **(A-D)** Concentrations of faecal cytokines (pg/mL) normalized to protein (ug/mL). **(E)** Calprotectin was measured in faecal supernatant by ELISA. **(A-E)** Data shown as mean ± S.D. **(F-H)** Concentration of antibodies (ng/mL) in faecal supernatant. **(I)** Representative contour plots of gating strategy for luminal bacteria and IgA^+^IgK^+^ and IgG^+^IgK^+^ bacteria from representative samples and (**J**) associated quantification. **(K)** Flow cytometry plots from lamina propria fractions of colon and associated quantification of B cells (%CD19^+^ of CD45^+^). If data was not normally distributed by Shapiro-Wilk it was log-transformed. Data analysed by one-way ANOVA with Dunnett’s multiple comparisons test, comparison to naive. N=8-9 per group **(A-H)**, n=6-8 per group **(J-K)**. Data shown as mean ± S.D. Dotted line is the mean of the naïve group.

Acute loss of intestinal immune cells, specifically in the Peyer’s Patches (PP), has previously been described after stroke (Brea et al., 2021; Hurry et al., 2024; Schulte-Herbrüggen et al., 2009; Tuz et al., 2024). In agreement, we saw reduced PP cellularity, driven by a loss of CD4 T cells and B cells at 5 days post stroke (Fig. 4E**-H** **and Fig S3B**) and the proportion of intraepithelial lymphocytes remained unchanged (**Fig. S3C-F**). This suggests reduced luminal cytokines may be driven by a loss of lymphocytes in the PP acutely after stroke. In contrast, the level of faecal calprotectin, a proinflammatory factor produced by neutrophils and monocytes that plays an important role in maintaining intestinal homeostasis, was reduced at two weeks and three months post-stroke (Fig. 4I), indicating distinct alterations to the immune environment during chronic recovery. The concentration of luminal IgA, the most abundant antibody isotype in the gut, remained unchanged following stroke througought the recovery period (Fig. 4J). Similarly, IgM levels showed no significant changes (Fig. 4K). In contrast, we observed a notable increase in IgG concentrations in the faecal supernatant three months post-stroke (Fig. 4L), driven primarily by increases in IgG1, IgG2b and IgG3 isotypes (**Fig. S4A**).

Alterations in the antibody coating of intestinal bacteria have been linked to post-stroke dysbiosis (Hurry et al., 2024; Schulte-Herbrüggen et al., 2009) acutely after stroke. Next, to understand if the increased IgG in the intestinal lumen observed a 3 months post stroke altered the antibody coating of intestinal bacteria, flow cytometry was performed on bacteria isolated from faecal pellets (Fig. 4M). Surprisingly, at three months post-stroke, the timepoint at which faecal IgG was elevated, bacteria showed less surface coating of both IgA and IgG (Fig. 4N). Reduced antibody coating was also seen on bacteria isolated from sham-operated animals, and as these animals were co-housed with stroked cagemates and exhibited the same gut microbiome composition as the stroke animals (**Fig. S4C**), this suggests that the type of bacteria present may be driving the reduced bacterial coating, rather than the availability of antibody. Indeed, a trend towards increased luminal IgG concentration was seen in sham-operated animals at three months (Fig. 4L). Furthermore, no increase in lamina propria B cells, the major source of antibody secreted into the intestinal lumen, was observed (Fig. 4O).

These results highlight that the immunological environment in the intestine throughout stroke recovery is altered during both acute and chronic recovery. The coexistence of increased unbound IgG and increased bacteria-bound IgG suggests potential interplay between antibody binding and availbility and bacterial community composition during chronic stroke recovery.

## 4. DISCUSSION

Post-stroke intestinal dysfunction has been extensively characterised during acute recovery and a relationship between the intestinal environment and the recovering stroke brain is clear. Here, we build on these findings with a longitudinal analysis that demonstrates the changes to intestinal homeostasis that take place over the course of recovery. To our knowledge this is the first description of intestinal alterations and microbial community chages during chronic experimental stroke recovery. Importantly, these changes are dynamic over time in the intestine, with the early resolution of functional deficits, but both the bacterial community composition and soluble immune factors displaying distinct profiles during acute and chronic phases of recovery. Although causation cannot be inferred from the current study, we provide some insight into possible interplay between immunological factors and microbial community composition in the intestine throughout stroke recovery.

Histological analyses were focused on the colon as this carries the highest bacterial burden across the intestine (Sekirov et al., 2010) and a shortened colon can be used as a proxy measure of colonic inflammation (Liang et al., 2024; Seo et al., 2024). We observed blunted crypt depth with reduced crypt cellularity throughout the recovery timecourse. Epithelial cell death, induced by activation of sympathetic nerves, has been reported to occur in both the colon and small intestine in hyperacute recovery from experimental stroke (Prame Kumar et al., 2023a) and combined with the trend for reduced proliferation we observed during acute recovery, may contribute to these changes. However, factors that contribute to prolonged changes to colon morphology are yet to be elucidated. Changes to the functional capacity of the gut have also been widely described during hyperacute recovery from stroke. We show that gut barrier function is intact at day five and remains intact therein, in agreement with published data showing integrity restoration as early as 24 hours post stroke (Hurry et al., 2024; Prame Kumar et al., 2023a; Stanley et al., 2016). We found gut transit time remained impaired at our acute five day timepoint but was restored in chronic recovery. Recent studies have shown that impaired peristalsis is driven by altered neural output in the gut after stroke (Kumar et al., 2024) and reduced mucin production may also play a role in delayed transit times (Blasco et al., 2020; Fang et al., 2023). In our studies, reduced food intake was observed alongside reduced mucin production and may further contribute to a prolonged transit time. As functional impairments and reduced food intake are confined to the acute recovery period it is plausible that these are important factors in the distinct microbiota alterations observed at this time point, although further studies would be required to confirm this.

Firmicutes and Bacteroidetes are the two most dominant phyla in both humans (Qin et al., 2010) and mice (Beresford-Jones et al., 2022) and the ratio of their abundance (F:B ratio) has been assessed in association with several pathological conditions including obesity (LEY Nat 2006), type 2 diabetes (Kochumon et al., 2024) and cardiovascular disease (Tsai et al., 2021). Alterd F:B ratio also occurs acutely after experimental stroke (Spychala et al., 2018) which we have shown increase in magnitude of change as recovery progresses. Firmicutes is reported to demonstrate reduced antibody surface coating than other enteric flora phyla (Bunker et al., 2015; Planer et al., 2016) and the expansion of intestinal firmicutes over time may drive the overall reduction in bacterial antibody coating found at chronic time points, independently of antibody availability. We chose to co-house the stroked animals with controls, in line with the IMPROVE Guidelines (Ischaemia Models: Procedural Refinements Of in Vivo Experiments), to enhance their welfare throughout the chronic recovery experiments (Percie du Sert et al., 2017). Mice are corporophagic and seeder mice can be used experimentally to modify the microbiome of their cagemates (Benakis et al., 2016). As a consequence of co-housing, we observed that the microbiome of the sham animals took on the profile of stroke animals, meaning that sham-induced alterations in intestinal homeostasis from naïve controls may not only be attributable to the surgical procedure. This experimental set up can aid hypothesis generation about cause and consequence when comparing dysbiosis alongside changes to the intestinal microenvironment. For example, changes to luminal IgG2b are significantly increased in both sham and stroked animals, and trend above baseline in other IgG isotypes, suggesting that dysbiosis may in part be driving changes to luminal antibody profiles. Together, these associations should allow future hypothesis-driven research to understand the precise mechanisms of gastrointestinal disturbances, with a particular focus on antibody coating of bacteria, during chronic stroke recovery and pave the way for potential therapeutic intervention.

A caveat of this study is that data were aquired from young, healthy, male mice and factors such as age, sex and co-morbidities are not accounted for. Indeed, studies using aged mice have shown that intestinal inflammation and barrier dysfunction are exacerbated with age (Wen et al., 2019). In addition, there are discrepancies in the literature on whether goblet cells and mucin production increase or decrease after stroke (Prame Kumar et al., 2023a). Time of sampling post-stroke may in part account for this, however sex differences may play an important role as increased mucus production out to seven days after stroke was found to be specific to females, with males instead increasing their production of anti-microbial peptides (Lee et al., 2023). Future studies looking at chronic intestinal disturbance in aged, mixed sex and co-morbid mice are required to expand on the initial insights we provide, however, as these factors can independently influence the intestinal environment, it is important to know the impact stroke, which itself imparts a complex mix of signals on the intestine, prior to adding complicating factors. Despite the need for further studies, our initial data suggests that stroke results in persistent changes in gut physiology along with altered antibody availability and microbial composition.

## 5. CONCLUSIONS

This study provides foundational understanding into the evolution of gastrointestinal disturbances after stroke. The data presented here demonstrate that experimental stroke induces significant alterations in the intestinal environment, with these changes evolving dynamically across time and peristing into the chronic recovery phase. Here, we demonstrate that the intestinal microbiota and immunoloigical profile is altered during chronic recovery, though whether these confer vulnerability or protection to additional insults, such as enteric infection, or modulate subsequent neurological insults, such as recurrent stroke, is unclear. Further studies will be required to determine whether persistent changes to these microbial families have a direct role in conferring poorer stroke outcomes throughout recovery.

## Author contributions

Concept: LM, ICM. Research Design: LM, ICM, RMLM, Data collection and analysis: LM, ICM, RMLM, RW, LMH, AM. Interpretation: LM, ICM, RMLM, RW, LMH, AM, CJA, DD, CCB, G-TH. Preparation of figures: LM, ICM, RMLM, RW. Manuscript writing: LM, ICM, RMLM. Manuscript editing: LM, ICM, RMLM, RW, LMH, AM, CJA, DD, CCB, G-TH.

## CRediT authorship contribution statement

**Rachel M. L. Martin** Data curation, formal analysis, investigation, methodology, visualisation, writing-original draft, writing-review and editing. **Isobel C. Mouat** Conceptualisation, Funding acquisition, Data curation, formal analysis, investigation, methodology, visualisation, writing-original draft, writing-review and editing. **Robert Whelan** Data curation, Formal analysis, methodology, visualisation, writing-review and editing. **Lizi M. Hegarty** Data curation, investigation, writing-review and editing. **Christopher J. Anderson** Writing-reviewing and editing **David H. Dockrell** Supervision, Writing-reviewing and editing **Calum C. Bain** Writing-reviewing and editing **Gwo-Tzer Ho** Writing-reviewing and editing **Laura McCulloch** Conceptualisation, Funding acquisition, Data curation, formal analysis, investigation, methodology, project administration, supervision, visualisation, writing-original draft, writing-review and editing.

## Acknowledgments

We appreciate the contributions of the NU-OMICS DNA Sequencing facility for carrying out the 16S rRNA sequencing. We gratefully acknowledge the support of the IRR core facilities including the IRR Flow Cytometry Facility and the IRR SURF Molecular Histology Microscopy. This work could not be completed without the support of our animal technicians in Bioresearch and Veterinary Services LF2 facility. We also thank Dr Frӓnze Progatzky for advice on gut transit time experiments.

This work was supported by a Wellcome Trust Sir Henry Dale Fellowship jointly funded by the Wellcome Trust and Royal Society grant number 220755/Z/20/Z and by the UKRI Biotechnology and Biological Sciences Research Council (BBSRC) grant number BB/T00875X/1. C.J.A. is supported by Wellcome Trust Career Development Award (225923/Z/22/Z). C.C.B. receives funding from UKRI/MRC (MR/W004763/1, MR/X018733/1) and G-T.H. receives funding from Helmsley Charitable Trust (G-1911-03343)

## Data availability

16S rRNA sequencing data that supports findings of this study are deposited in Edinburgh DataShare with the accession number https://doi.org/10.7488/ds/7922. All other data supporting the findings of this study are available from corresponding author upon reasonable request.

## Open Access statement

This research was jointly funded in whole or in part by the Wellcome Trust and the Royal Society [Grant Number 220755/Z/20/Z]. For the purpose of open access, the author has applied a CC-BY public copyright licence to any author accepted manuscript version arising from this submission.

---

**Figure S1:**
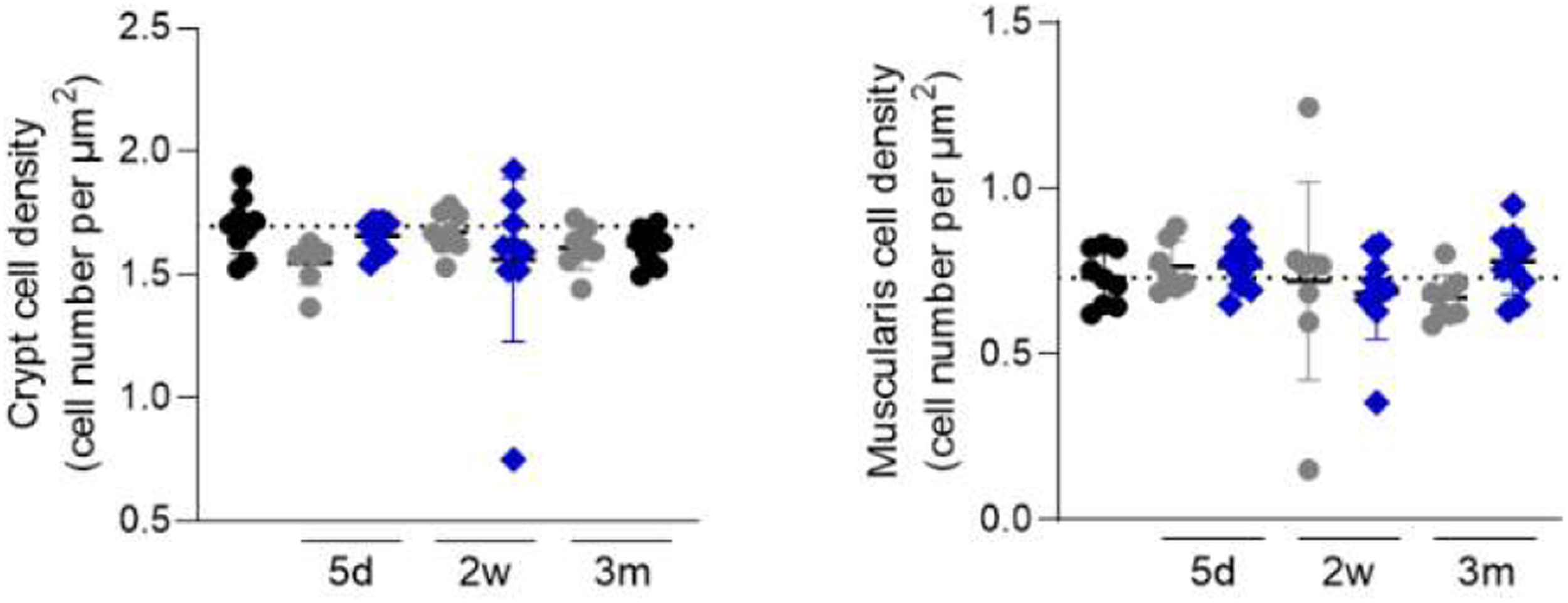
Effect of experimental stroke on crypt and muscularis cell density. The number of nuclei quantified per µm^2^ in the colonic crypt and muscularis. Data shown as mean ± S.D. Dotted line is the mean of the naïve group. Data analysed by one-way ANOVA with Dunnett’s multiple comparisons test, comparison to naive. N=7-10 per group.

**Figure S2:**
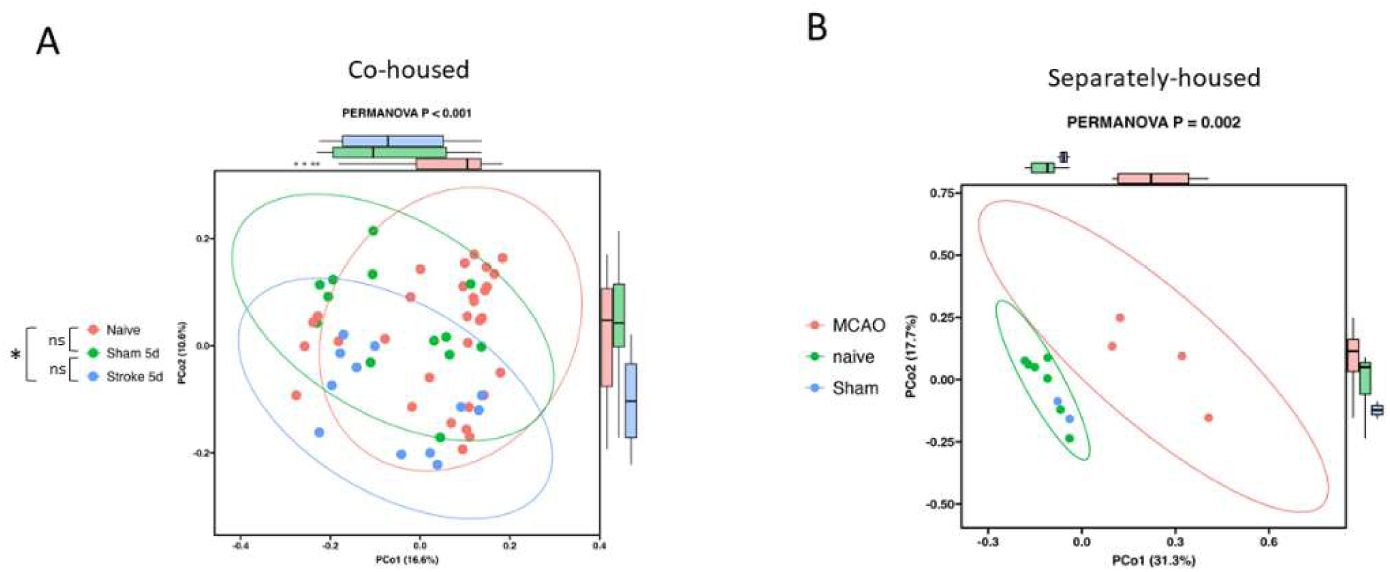
Co-housing impacts the microbial composition of controls. Bray-Curtis dissimilarity index analysis of **(A)** sham and stroke animals co-housed, naïve animals housed separately and **(B)** naïve, sham, and stroke animals all separately housed. **(A)** Data compiled from two independent experiments, n=12-25. **(B)** N=2-7. Analysed by PERMANOVA, with and without multiple comparisons test.

**Figure S3:**
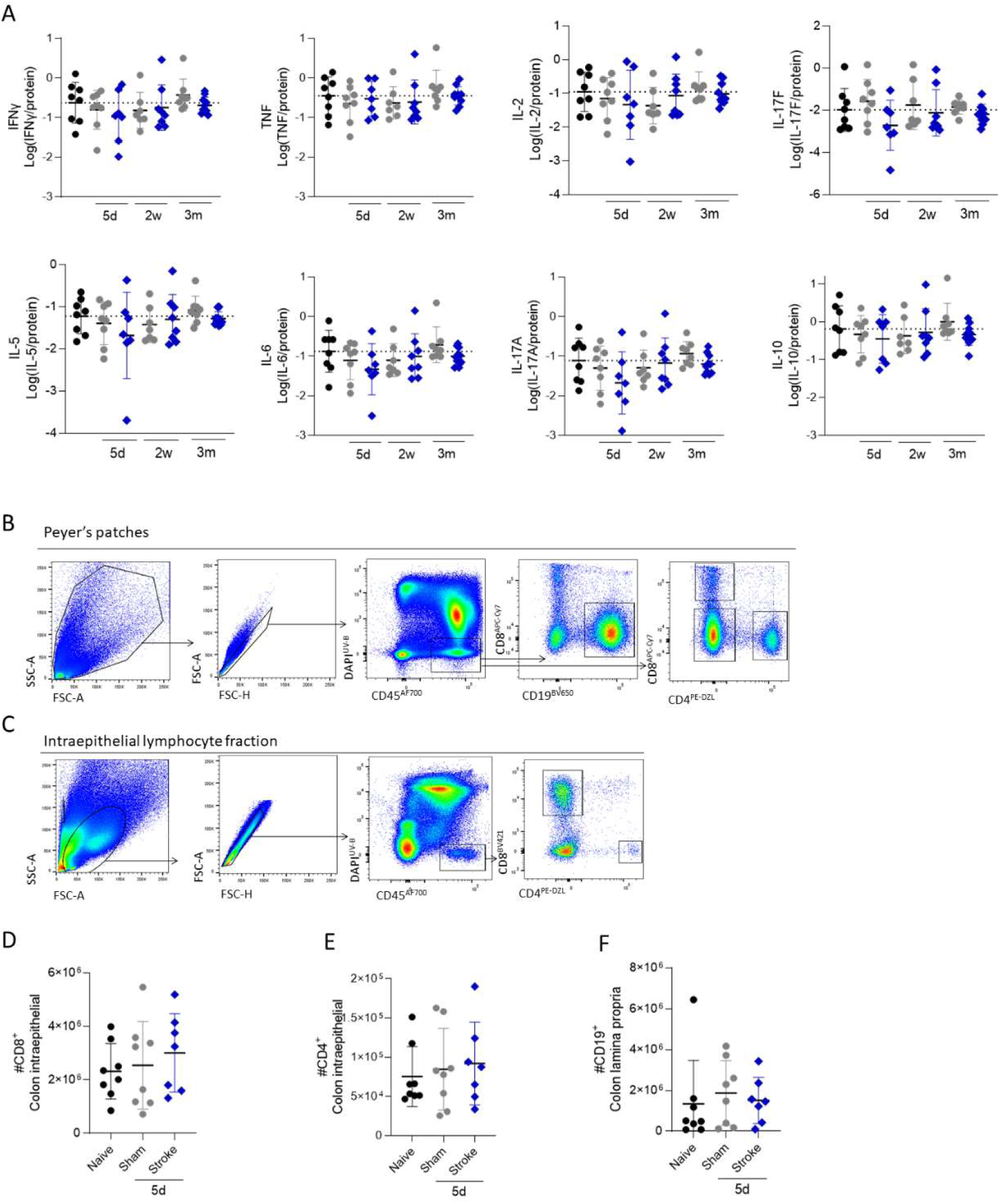
The abundance of various cytokines in faecal supernatant are unchanged over experimental stroke recovery. **(A)** Concentrations of faecal cytokines (pg/ml) normalized to protein (ug/mL). Data shown as mean ± SD. Data was not normally distributed by Shapiro-Wilk and therefore was log­transformed. Log-transformed data was analysed by one-way ANVOA with comparison to naive. Dotted line is the mean of the naive group. N=7-10 per group. **(B)** Peyer^1^s patch cell number. N=7-8 per group, analysed by one-way ANVOA with comparison to naïve.

**Figure S4:**
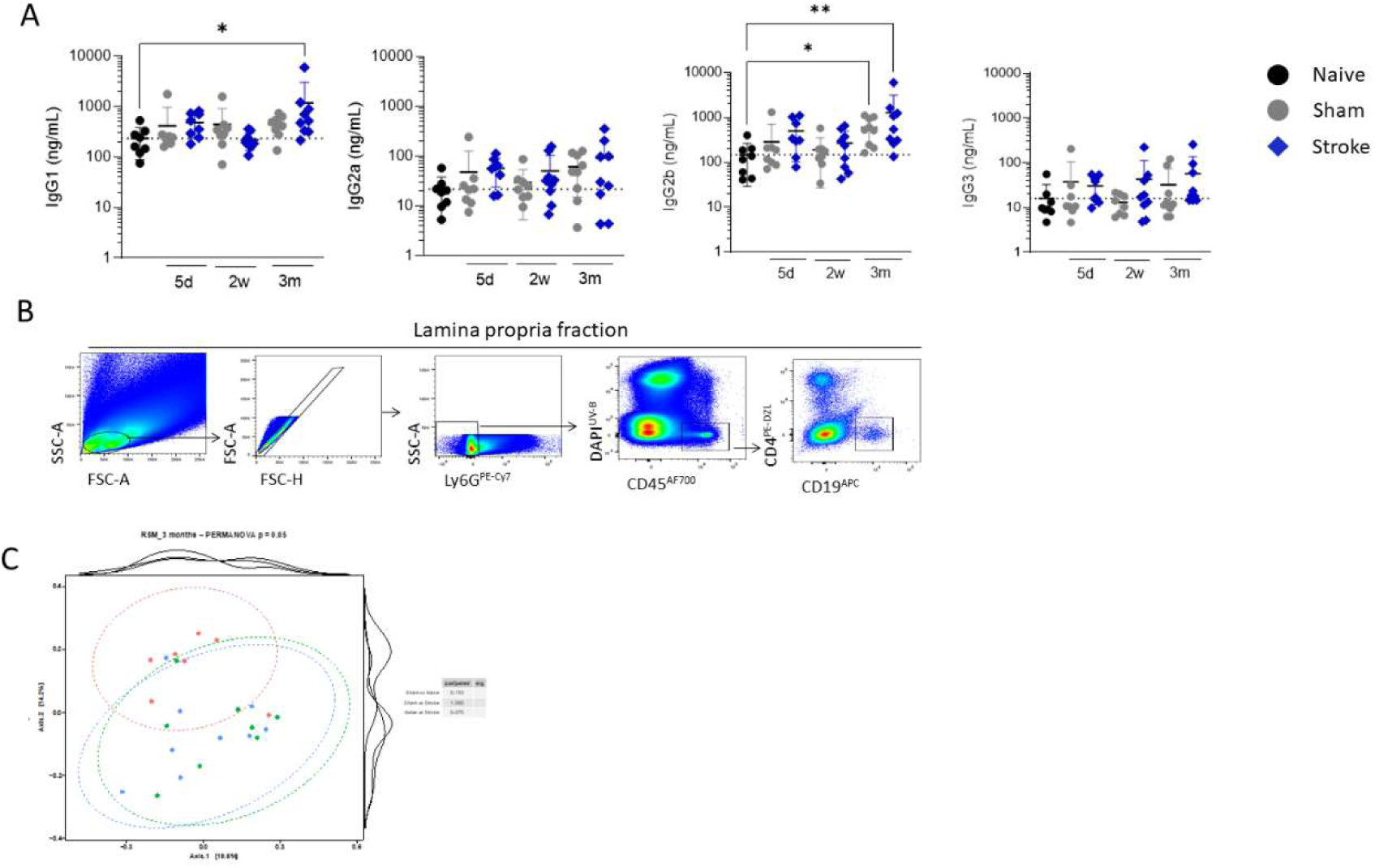
Antibodies subclasses, Peyer’s patch gating scheme. **(A)** Concentration of IgG subclasses (ng/mL) in faecal supernatant. **(B)** Gating scheme for lymphocytes in the lamina propria of colons. **(C)** PCA plots of intestinal bacterial community composition, measured by 165 rDNA sequencing; sham and stroke animals co-housed, naïve animals separately housed. **(A)** Data was analysed by one-way ANOVA, or Kruskal-Wallis if not normally distributed, with multiple comparisons with comparison to naïve. Dotted line is the mean of the naïve group. **(C)** PERMANOVA, with and without multiple comparisons test. **(A)** N=7-10 per group **(C)** N=7-10 per group.

